## Supplementary Figures for "Nucleocytoplasmic transport of active HER2 causes fractional escape from the DCIS-like state"

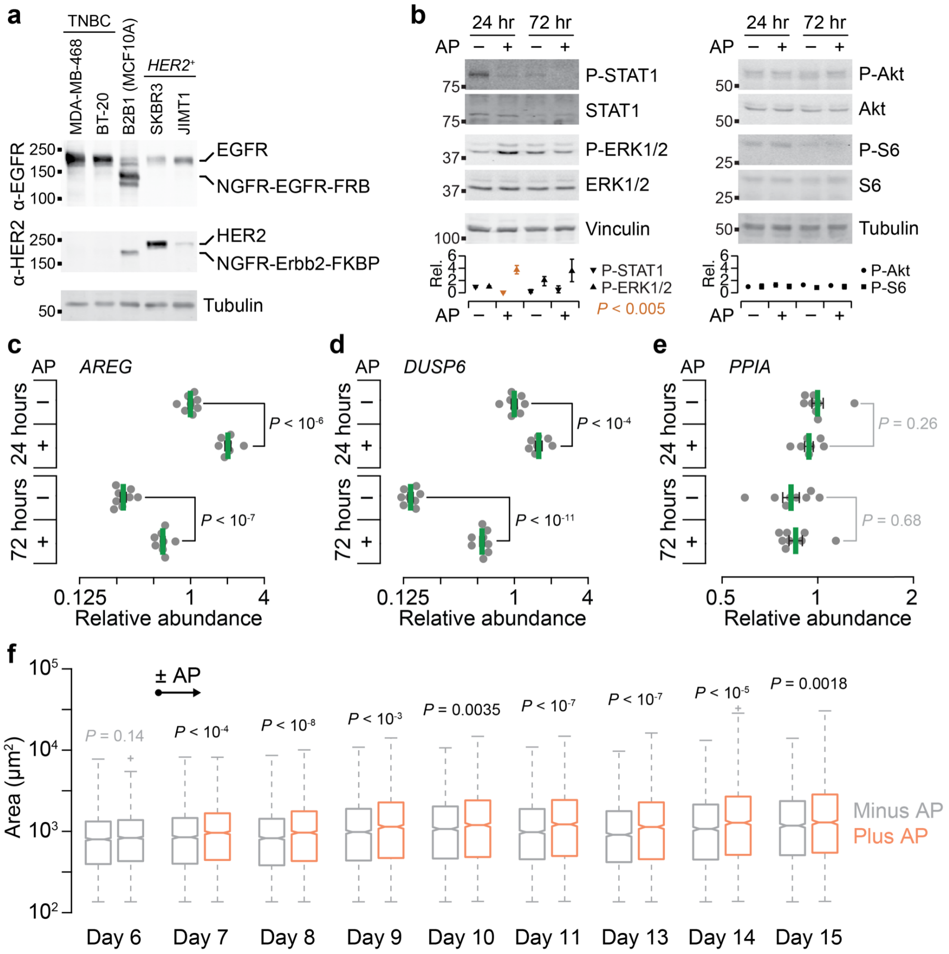


**Supplementary Fig. 1 | A B2B1 subclone of MCF10A-5E cells with moderate ectopic expression of chemically activatable EGFR and Erbb2 chimeras.**

**a**, Abundance of NGFR-EGFR-FRB and NGFR-Erbb2-FKBP chimeras in B2B1 cells is comparable to endogenous expression in triple-negative breast cancer (TNBC) and *HER2*-amplified (*HER2*$^{+}$) breast cancer lines respectively. Protein extracts were immunoblotted for EGFR and HER2 intracellular domains with tubulin used as a loading control.

**b**, ErbB heterodimerization alters phosphorylation of STAT1 (Tyr701) and ERK1/2 (Thr202/Tyr204) but not phosphorylation of Akt (Ser473) or S6 (Ser240/244). Protein extracts from 3D-cultured B2B1 cells treated ±0.5 μM AP for the indicated times after Day 6 were immunoblotted for the indicated proteins with total STAT1, total ERK1/2, total Akt, total S6, vinculin, and tubulin used as loading controls. Bottom graphs show the mean ± range of replicated densitometry from $n$ = 3 biological replicates; at each time point, differences between ±AP conditions were assessed by two-sided $t$ test.

**c–e**, ErbB heterodimerization induces canonical ErbB target genes with no detectable effect on housekeeping genes. Quantitative PCR for *AREG* (**c**), *DUSP6* (**d**), and *PPIA* (**e**) in B2B1 cells cultured in 3D for six days ±0.5 μM AP21967 (AP) for 24 hours or 72 hours where indicated. Data are shown as the geometric mean (normalized to the 24-hour, minus-AP condition) ± log-transformed s.e. from $n$ = 7–8 biological replicates. Differences in geometric means were assessed by two-sided $t$ test after log transformation.

**f**, ErbB heterodimerization causes a global and sustained increase in outgrowth size within 24 hours of AP addition. B2B1 cells were cultured in 3D for six days followed by stimulation with or without 0.5 μM AP for the indicated times after Day 6. Cross-sectional areas were quantified for $n$ = 1851–2229 (Minus AP) or 1774–2317 (Plus AP) outgrowths from four biological replicates. Boxplots show the median area, estimated 95% confidence interval of the median (notches), interquartile range (box), 1.5x the interquartile range from the box edge (whiskers), and outliers (+). Differences between groups for each time point were assessed by two-sided rank sum test with Šidák correction for multiple-hypothesis testing.

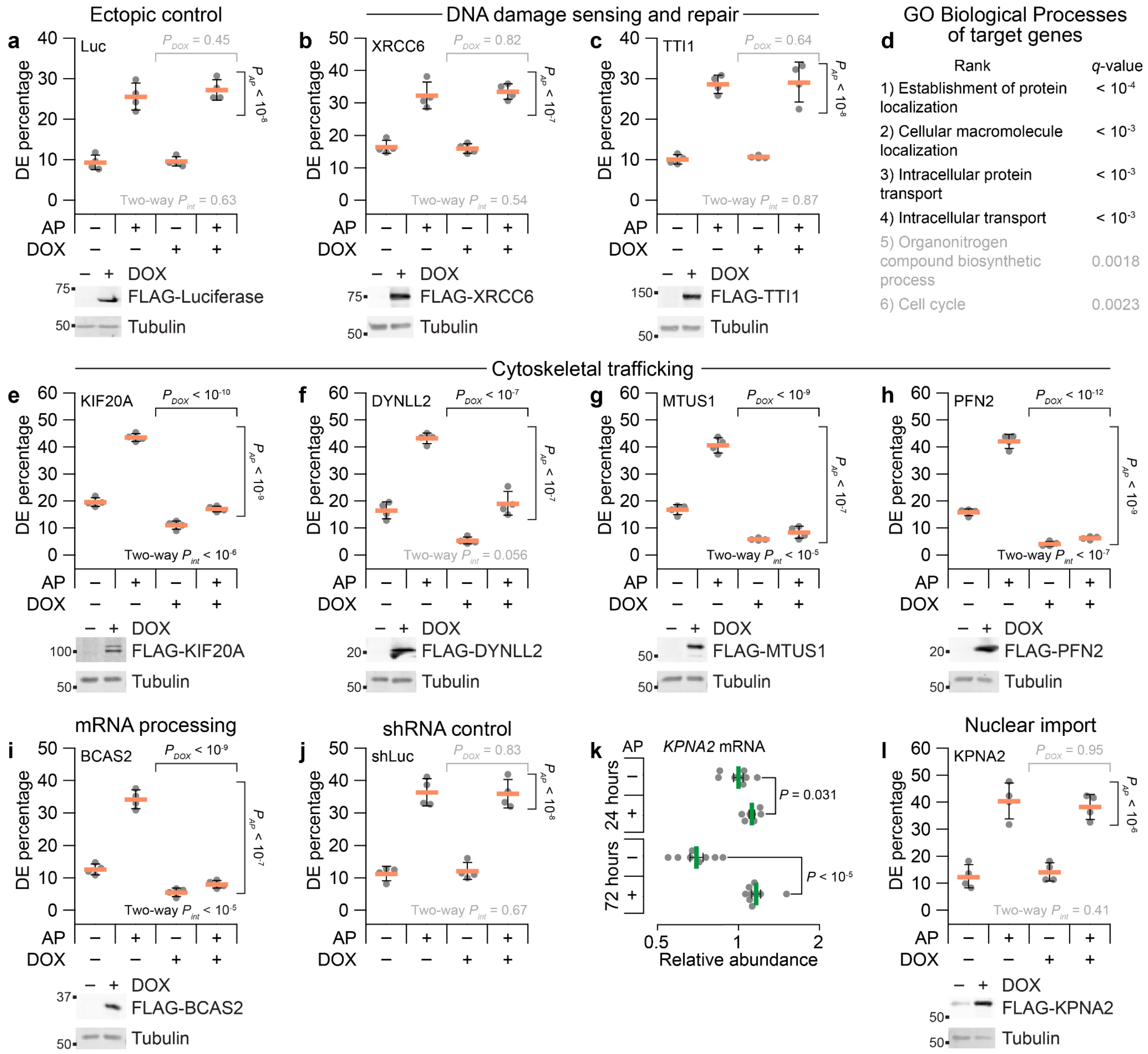


**Supplementary Fig. 2 | Exclusion of alternative hypotheses about other candidates.**

**a**, DE percentage for B2B1 cells stably expressing inducible FLAG-tagged luciferase.

**b**, **c**, Candidates related to DNA damage sensing–repair do not affect DE penetrance. B2B1 cells expressing inducible FLAG-tagged XRCC6 (**c**) or TTI1 (**d**) were used where indicated.

**d**, Enrichment analysis of gene ontology (GO) biological processes for the 97 candidate genes (= 15 high-priority targets + 86 candidates – [4 uncharacterized loci and probeset redundancies]) identified by stochastic frequency matching. False discovery rate-corrected *q*-values are shown for each biological process.

**e**–**h**, Candidates related to cytoskeletal trafficking inhibit ErbB-induced DE penetrance. B2B1 cells expressing inducible FLAG-tagged KIF20A (**e**), DYNLL2 (**f**), MTUS1 (**g**), or PFN2 (**h**) were used where indicated.

**i**, A candidate amplified in breast cancer[1] and involved in mRNA processing inhibits ErbB-induced DE penetrance. B2B1 cells expressing inducible FLAG-tagged BCAS2 were used.

**j**, DE percentage for B2B1 cells stably expressing inducible shRNA targeting luciferase (shLuc) as an shRNA control.

**k**, Quantitative PCR for *KPNA2* in B2B1 cells cultured in 3D for six days followed by addition of 0.5 µM AP21967 (AP) for the indicated times. Data are shown as the geometric mean (normalized to the 24-hour, minus-AP condition) ± log-transformed s.e. from $n$ = 7–8 biological replicates. Differences in geometric means were assessed by two-sided $t$ test after log transformation.

**l**, A regulator of nuclear import does not affect ErbB-induced DE penetrance. B2B1 cells expressing inducible FLAG-tagged KPNA2 were used.

For **a–c**, **e–j**, and **l**, B2B1 cells stably expressing doxycycline (DOX)-inducible ectopic constructs were 3D cultured for 9–13 days with 0.5 μM AP added at Day 6 and/or 1 μg/ml DOX added at Day 5 where indicated. Data are shown as the arcsine transformed mean ± s.e. from $n$ = 4 biological replicates where >100 outgrowths were scored per replicate. Differences by factor (DOX or AP) and two-factor interaction (int) were assessed by two-way ANOVA after arcsine transformation. Ectopic expression was confirmed by immunoblotting for FLAG with tubulin used as a loading control.

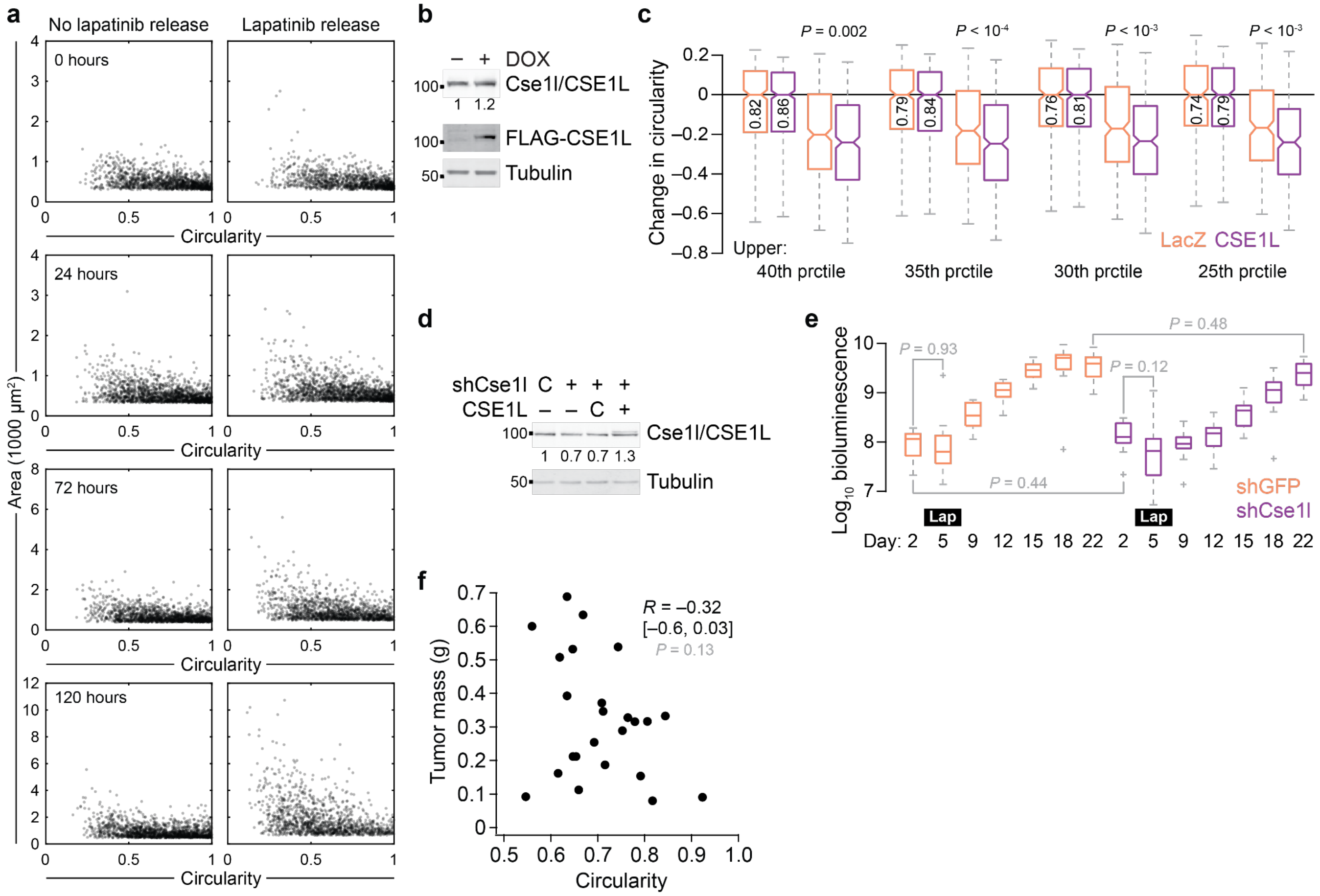

**Supplementary Fig. 3 | 3D and in vivo characterization of TM15c6 cells with or without pharmacologic and genetic perturbations.**

**a**, Changes in cross-sectional area upon lapatinib release do not noticeably skew changes in circularity. TM15c6 cells 3D cultured for five days in 2 µM lapatinib followed by release for the indicated time. Cross-sectional area and circularity were analyzed for all outgrowths ($n$ = 1241–1755 outgrowths from four biological replicates).

**b**, Inducible ectopic expression of CSE1L confirmed by immunoblotting for total Cse1l/CSE1L and FLAG with tubulin used as a loading control. Inducible overexpression of total Cse1l/CSE1L was quantified relative to the minus-DOX control.

**c**, CSE1L-induced circularity phenotype is not sensitive to the choice of cross-sectional area thresholding. Circularities were analyzed for outgrowths above the indicated threshold by size ($n$ = 504–1018 outgrowths from four biological replicates).

**d**, Inducible knockdown of Cse1l and addback of CSE1L. TM15c6 cells stably expressing inducible shCse1l or shGFP control (C) with or without inducible CSE1L or LacZ control (C) were treated with 1 µg/ml doxycycline for two days and immunoblotted for Cse1l/CSE1L with tubulin used as a loading control.

**e**, Dynamics of tumor bioluminescence in TM15c6 cells expressing shCse1l or shGFP control after inoculation at Day 0 and administration of 30 mg/kg lapatinib i.p. daily from Days 3–9 ($n$ = 12 animals).

For **c** and **e**, boxplots show the median bioluminescence, interquartile range (box), estimated 95% confidence interval of the median (**c**, notches), 1.5x the interquartile range from the box edge (whiskers), and outliers (+). Differences between indicated groups were assessed by two-sided rank sum test with Šidák correction for multiple-hypothesis testing.

**f**, Tumor size is not significantly correlated with tumor circularity. Mass and circularity of excised tumors ($n$ = 24 samples) were assessed by Pearson correlation ($R$) with 95% confidence interval and hypothesis testing for nonzero correlation estimated after Fisher Z transformation.

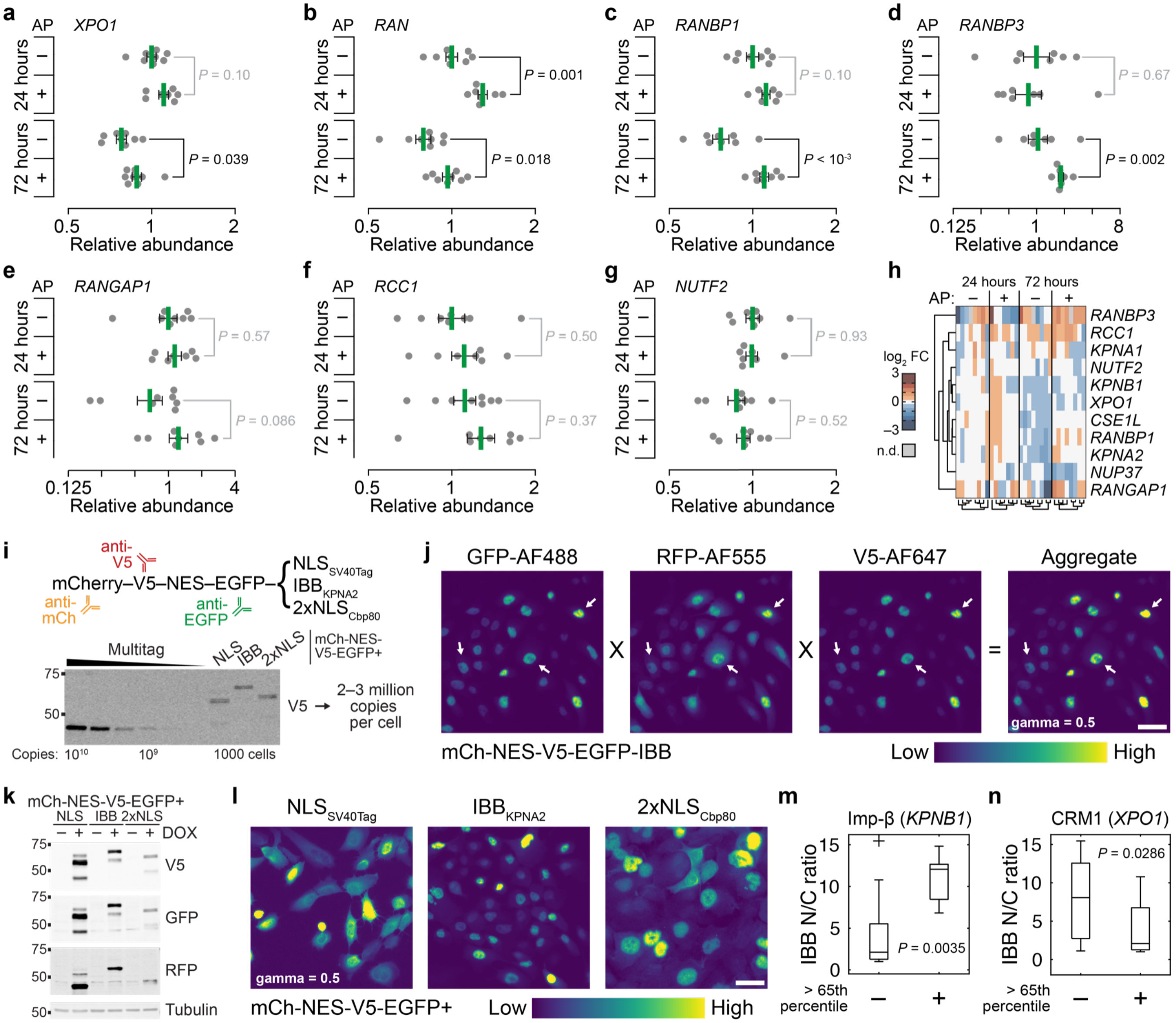


**Supplementary Fig. 4 | Motivation, testing, and use of a nucleocytoplasmic transport model.**
**a–g**, Quantitative PCR for *XPO1* (**a**), *RAN* (**b**), *RANBP1* (**c**), *RANBP3* (**d**), *RANGAP1* (**e**), *RCC1* (**f**), and *NUTF2* (**g**) in B2B1 cells cultured in 3D for six days followed by 0.5 µM AP21967 (AP) for 24 hours or 72 hours. Data are shown as the geometric mean (normalized to the 24-hour, minus-AP condition) ± log-transformed s.e. from $n$ = 7–8 biological replicates. Differences in geometric means were assessed by two-sided $t$ test after log transformation.

**h**, Replicate-by-replicate clustergram of the quantitative PCR data from Fig. 2g–j, Supplementary Fig. 2k, and Supplementary Fig. 4a–g. Data were normalized to the geometric mean of the 24-hour, minus-AP condition for each gene and row clustered (Euclidean distance, Ward's linkage). Replicate groups were column clustered separately

**i**, Tandem cargo reporters. mCherry (mCh), V5, and EGFP track steady-state accumulation of different NLSs paired with an NES. Absolute copy-number quantification was performed with recombinant V5-containing Multitag[2].

**j**, Image math for Alexa Fluor (AF)-based immunolocalization of GFP, RFP/mCh, and V5 to estimate full-length tandem cargo reporter.

**k**, Inducible expression of tandem cargo reporters. B2B1 cells expressing inducible mCh-NES-V5-EGFP tagged with NLS$_{SV40Tag}$, IBB$_{KPNA2}$, or 2xNLS$_{Cbp80}$ induced with 1 µg/ml doxycycline (DOX) for 6 hours were immunoblotted for the indicated epitopes with tubulin used as a loading control.

**l**, Representative image math immunolocalization of the indicated tandem cargo reporters after DOX induction for 6 hours. Quantification is summarized in Fig. 3b–d.

**m**, **n**, Proportionately high expression of Imp-β and CRM1 alters the steady-state accumulation of Imp-β-binding cargo. Predicted IBB nuclear/cytoplasmic (N/C) ratio for *n* = 20 AP-treated B2B1 outgrowths split at the 65th percentile according to relative transcript abundance of *KPNB1* (**e**), or *XPO1* (**f**). Boxplots show the median N/C ratio (horizontal line), interquartile range (box), 1.5x the interquartile range from the box edge (whiskers), and outliers (+). Differences between groups were assessed by a one-sided rank sum test.
For **j** and **l**, aggregate images are displayed with image gamma = 0.5, and the scale bar is 20 μm.

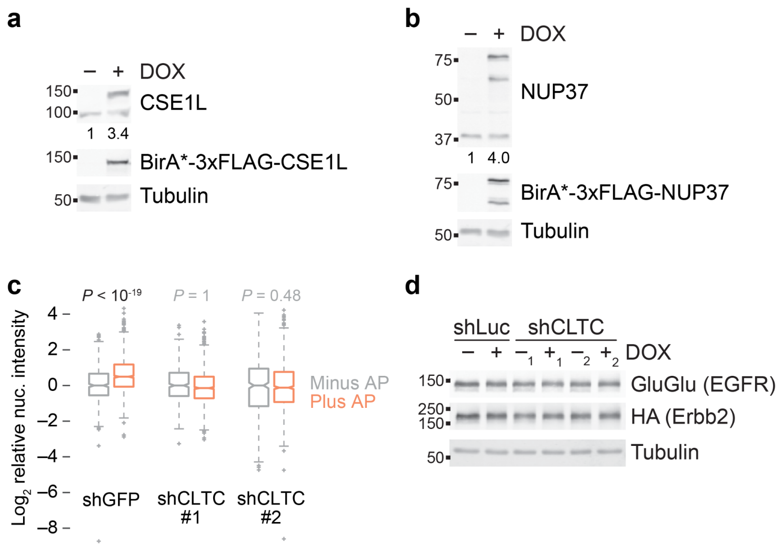


**Supplementary Fig. 5 | Extended data related to CSE1L–NUP37 proximity labeling and CLTC knockdown.**
**a, b**, Induction of ectopic BirA*-CSE1L (**a**) or BirA*-NUP37 (**b**) relative to endogenous protein. Inducible expression was compared relative to the no DOX control with tubulin used as a loading control.
**c**, CLTC knockdown inhibits the internalization of EGFR–Erbb2 heterocomplexes. B2B1 cells expressing the indicated shRNAs were plated on coverslips and induced with 1 μg/ml DOX for 48 hours before treatment with 0.5 μM AP for 15 minutes and immunostaining for GluGlu and HA tags. Boxplots show the normalized nuclear intensity (relative to minus AP), interquartile range (box), estimated 95% confidence interval of the median (notches), 1.5x the interquartile range from the box edge (whiskers), and outliers (+) from $n$ = 433–1146 cells of two or four biological replicates. Differences between AP groups were assessed by one-sided rank sum test.
**d**, CLTC knockdown does not alter long-term abundance of EGFR or Erbb2 chimeras. B2B1 cells expressing the indicated shRNAs were immunoblotted for GluGlu and HA tags with tubulin used as a loading control.

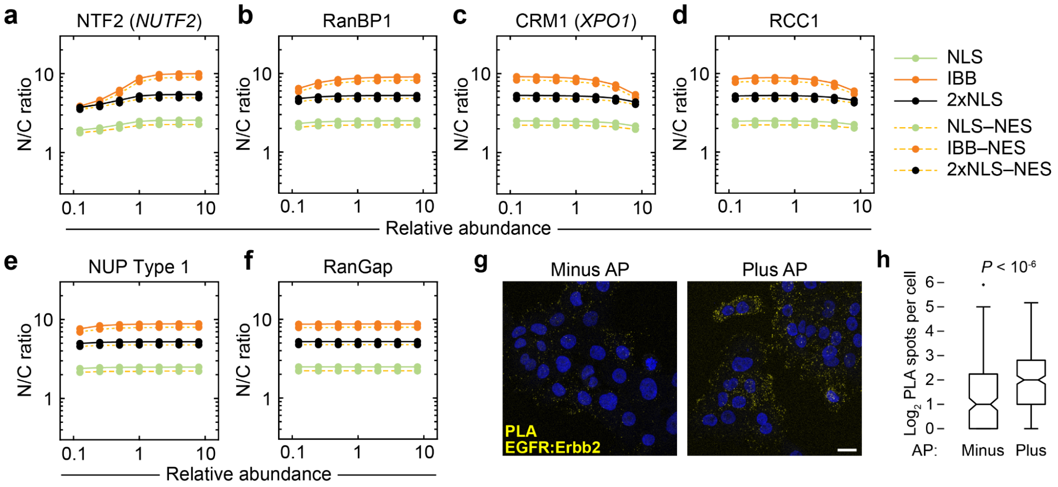


**Supplementary Fig. 6 | Extended sensitivity analysis of the model and validation of the proximity ligation assay.**

**a–f,** Predicted sensitivity of the systems model to starting protein concentrations of NTF2 (**a**), RanBP1 (**b**), CRM1 (**c**), RCC1 (**d**), total NUP (**e**), and RanGap (**f**). Protein abundances are scaled relative to the concentrations used in the base model (Supplementary Table 4). Steady-state nuclear-to-cytoplasmic (N/C) ratios are shown for 1 µM of the representative cargo described in Fig. 4a.

**g,** The proximity ligation assay detects regulated protein-protein interactions. B2B1 cells were treated with or without 0.5 µM AP for 15 minutes and costained for interaction between the EGFR and Erbb2 chimeras. Scale bar is 20 µm.

**h,** Quantification of PLA spots in B2B1 cells treated with or without 0.5 µM AP and stained as in (**g**). Boxplots show the median PLA spots per cell (horizontal line), interquartile range (box), estimated 95% confidence interval of the median (notches), 1.5x the interquartile range from the box edge (whiskers), and outliers (+) from $n$ = 195–272 cells from eight confocal image stacks on one coverslip per condition. Differences between groups were assessed by two-sided KS test with Bonferroni correction.

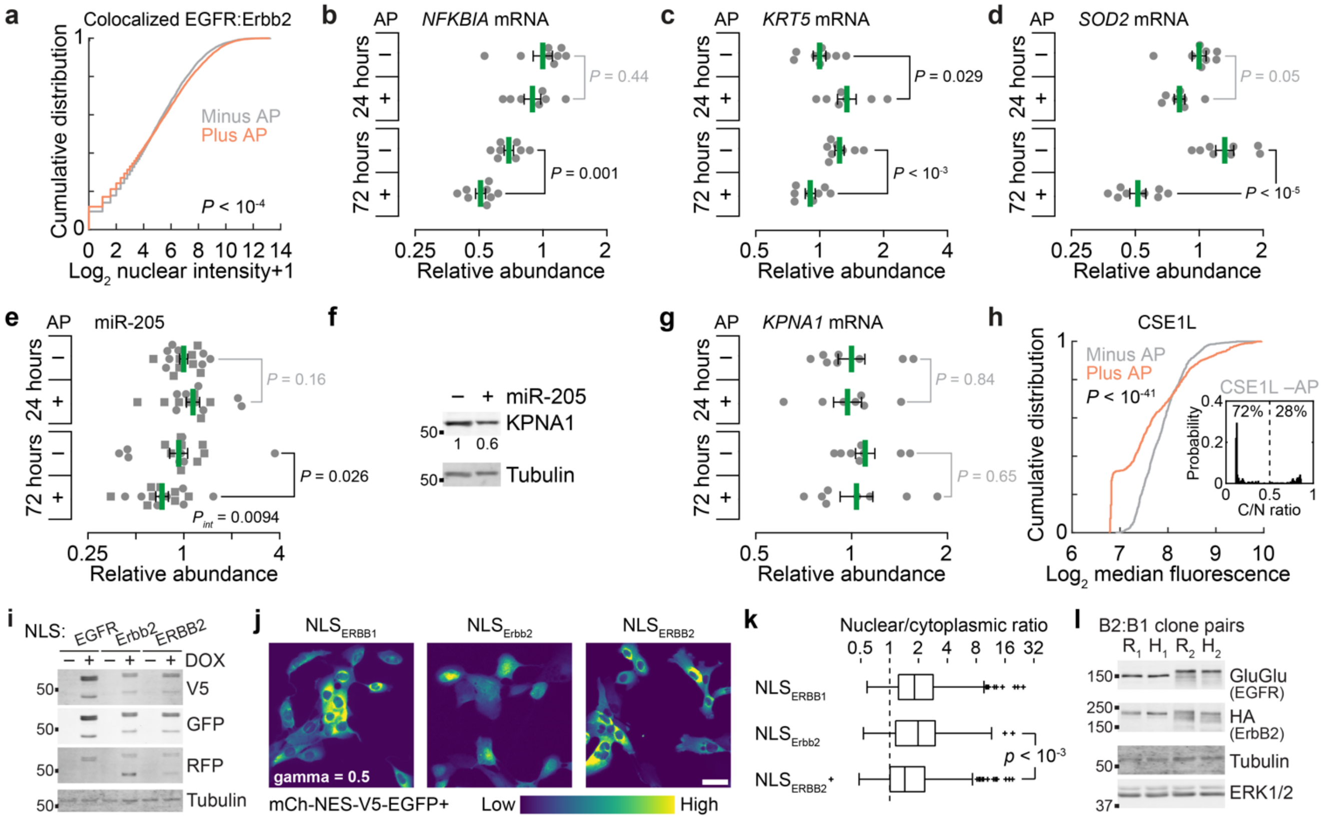

**Supplementary Fig. 7 | Genomic loci bound by chimeric EGFR, controls for miR-205, and comparisons of ErbBs from primates and rodents.**

**a**, Colocalized nuclear EGFR:Erbb2 staining in B2B1 cells cultured in 3D for six days ±0.5 μM AP for 24 hours. Differences were assessed by two-sided KS test.

**b–e**, Quantitative PCR for *NFKBIA* (**b**), *KRT5* (**c**), *SOD2* (**d**), and miR-205 (**e**) in B2B1 cells cultured in 3D for six days ±0.5 μM AP21967 (AP) for 24 hours or 72 hours.

**f,** miR-205 suppresses KPNA1. 293T cells were transfected with miR-205 precursor or negative control for 48 hours and immunoblotted for KPNA1 with tubulin used as a loading control.

**g,** Quantitative PCR for *KPNA1* in B2B1 cells cultured in 3D for six days ±0.5 μM AP21967 (AP) for 24 hours or 72 hours where indicated.

**h,** CSE1L quantification in B2B1 cells as in Fig. 7h. (Inset) Model predictions (Fig. 7k) when CSE1L is reduced to 40% (reflecting 3D culture; Fig. 2g) and randomly perturbed (**h**, gray). Simulations were performed as in Fig. 7k with $K_D$ = 3 nM, and steady-state cytoplasmic/nuclear (C/N) ratios above or below 50% are shown from *n* = 500 initializations.

**i**, Inducible ErbB tandem cargo reporters validated as in Supplementary Fig. 4k.

**j**,**k**, Representative immunolocalization (**j**) and quantification (**k**) after DOX induction for 6 hours. Boxplots show the median N/C ratio (horizontal line), interquartile range (box), 1.5x the interquartile range from the box edge (whiskers), and outliers (+) from *n* = 283 ($NLS_{ERBB1}$), 285 ($NLS_{Erbb2}$), and 333 ($NLS_{ERBB2}$) cells collected from four biological replicates. Differences were assessed by two-sided rank sum test. For **j**, the scale bar is 20 μm.

**l,** Characterization of ErbB clones expressing human ERBB2 or rat Erbb2. Protein extracts were immunoblotted for GluGlu and HA epitope tags with tubulin and ERK1/2 used as loading controls.

For **b–e** and **g**, data are shown as the geometric mean (normalized to the 24-hour, minus-AP condition) ± log-transformed s.e. from *n* = 7–8 biological replicates. Differences in geometric means were assessed by two-sided *t* test after log transformation.
