## Supplementary Note 1 for "Nucleocytoplasmic transport of active HER2 causes fractional escape from the DCIS-like state"

**Supplementary Note 1.** Development of the nucleocytoplasmic transport model.

*Model Reconstruction*

The nucleocytoplasmic transport model was based upon two earlier models of Riddick and Macara[1, 2]. The first model[1] encodes importin-α/β shuttling dynamics, the import of cargo harboring different types of nuclear localization sequence (NLS), and the transport of Ran. Code for the first model was obtained from the original publication as a Jarnac script[3], which we converted to MATLAB language with a Jarnac-to-MATLAB translator from the Systems Biology Workbench[4]. This model recreated most of the simulations reported in the publication[1] but required multiple edits to reflect accurately the experimental parameters drawn from the literature (see Model Corrigenda below).

The second model[2] builds upon the first by adding the machinery for nuclear export along with a dedicated subcompartment for nuclear pores. Regrettably, code for the second model could not be recovered from the BioModels database[5] where it was purportedly deposited[2]. As an alternative, we recoded the second model to the extent possible given the descriptions and parameters reported in Ref. [2]. The recoding required some assumptions that are elaborated upon here (see Nuclear Pore Complex, Nuclear Export Module, and Model Addenda below). The final model reflecting all changes and additions has been deposited in the BioModels database (MODEL2210060001) and is available on GitHub (JanesLab/NucCytoShuttle).

*Model Corrigenda*

Although the MATLAB translation of the Ref. [1] model was accurate, we uncovered multiple inconsistencies in reported parameter values after checking the original sources. Some were errors in transcription from the literature. For example, one cited study[6] quantifying the interaction between RanGTP and importin-β reports a half-life ($t_{1/2}$) of four hours for the RanGTP:importin-β complex. By definition, $t_{1/2}$ is related to the dissociation rate constant ($k_{off}$) as follows:

$$k_{off} = \frac{\ln(2)}{t_{1/2}} \tag{1}$$

yielding $k_{off}$ = 4.8 x $10^{-5}$ $s^{-1}$ for the RanGTP:importin-β complex. However, in Ref. [1] and in the original Jarnac code, this dissociation rate constant is listed as 4.8 x $10^{-6}$ $s^{-1}$, giving rise to a tenfold higher affinity in the model than that reported in Ref. [6]. In the same table of Ref. [1], association rate constants ($k_{on}$) are listed in units of $\mu M^{-1}s^{-1}$ but the values in the Jarnac code support units of $M^{-1}s^{-1}$, which are also more biophysically realistic. Likewise, bimolecular-to-biomolecular exchange reactions—such as importin-α–importin-β–NLS + RanGTP ↔ importin-α–NLS + importin-β–RanGTP—should have reverse rates in units of $M^{-1}s^{-1}$, not $s^{-1}$ as listed[1]. We corrected these parameters and others with evidence of transcription errors (Supplementary Table 1).

Elsewhere, we discovered inconsistencies in the reporting of model parameters. Ref. [1] lists parameters for the RanGAP-catalyzed hydrolysis of a RanGTP–CAS–importin-α–RanBP1 species, but this species does not exist in the Jarnac code. Instead, RanBP1 binding to RanGTP–CAS–importin-α converts to RanBP1–RanGTP + CAS + importin-α, which we retain in the final model. Estimates of other transport receptors and native cargo for importin-α–importin-β and importin-β are listed as whole-cell concentrations in Ref. [1] but used as cytoplasmic concentrations in the Jarnac code. We use the Ref. [1] numbers as whole-cell concentrations to stay consistent with the other proteins in the same table. The accumulating corrections and changes prompted us to abandon attempts at retaining all the modeling results of the earlier work.

Most problematic was the overall handling of interactions (importin-α–importin-β, importin-α–importin-β–NLS, and importin-β–cargo from Ref. [1]) that were modeled using two-state kinetics estimated from surface-plasmon resonance data of Catimel et al.[7]. A two-state model casts the bimolecular interaction of *A* and *B* as:

$$A + B \underset{k_{off,1}}{\overset{k_{on,1}}{\rightleftarrows}} AB^* \underset{k_{off,2}}{\overset{k_{on,2}}{\rightleftarrows}} A{:}B \tag{2}$$

where *AB** is a precursor complex, and *A:B* is the mature complex. Unlike a one-state association rate constant, $k_{on,2}$ is defined in units of per time. Refs. [1, 7] both list $k_{on,2}$ in units of per concentration per time. The difference is crucial because of the way that an effective dissociation constant ($K_{D,eff}$) is defined for a two-state model:

$$K_{D,eff} = \frac{k_{off,1}}{k_{on,1}\left(1+\frac{k_{on,2}}{k_{off,2}}\right)} \tag{3}$$

Ref. [7] lists the four rate parameters of the two-state model along with an estimated $K_{D,eff}$, which according to Equation 3 is incorrect by roughly one million-fold (~fM affinity instead of ~nM). The parameter table in Ref. [7] lists $k_{on,2}$ in units of $10^3$ $M^{-1}s^{-1}$—based on one text reference to the $k_{on,2}$ for the importin-β:cargo complex, we believe all of the $k_{on,2}$ values are instead in units of $10^{-3}$ $s^{-1}$. Changing the units in this way reconciles the two-state kinetic parameters of Ref. [7] with the accompanying estimate of $K_{D,eff}$ and implies that all $k_{on,2}$ parameters in Ref. [1] are incorrect by six orders of magnitude (Table SN1). The final model uses the revised set of parameters that are self-consistent with the original data source[7] (Supplementary Table 1).

*Nuclear Pore Complex*

To model nuclear pore complexes (NPCs), we began by creating a perinuclear subcompartment comprised of volume removed from the nuclear and cytoplasmic subcompartments. The latest cryoelectron microscopy data on the human NPC suggest that it is 70 nm tall[8, 9]. Using this dimension ($h_{NPC}$) and the volume of the nucleus specified in the model ($V_{nuc}$), we approximated the nucleus as a sphere and removed radial volume from the nucleus and cytoplasm to create the perinuclear volume ($V_{pn}$):

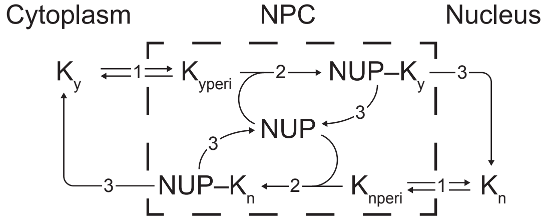


**Fig. SN1.** Modeling architecture for the transport of karyopherins (K) and karyopherin-containing complexes across the NPC subcompartment through a nuclear pore (NUP).

$$V_{pn} = \frac{4\pi}{3}\left[\left(\sqrt[3]{\frac{3}{4\pi}V_{nuc}} + \frac{h_{NPC}}{2}\right)^3 - \left(\sqrt[3]{\frac{3}{4\pi}V_{nuc}} - \frac{h_{NPC}}{2}\right)^3\right] \tag{4}$$

We then updated the volumes for the nucleus ($V_{nuc\text{-}pc}$) and cytoplasm ($V_{cyto\text{-}pc}$) accordingly:

$$V_{nuc-pc} = \frac{4\pi}{3}\left(\sqrt[3]{\frac{3}{4\pi}V_{nuc}} - \frac{h_{NPC}}{2}\right)^3 \tag{5}$$

$$V_{cyto-pc} = V_{cyto} + V_{nuc} - \frac{4\pi}{3}\left(\sqrt[3]{\frac{3}{4\pi}V_{nuc}} + \frac{h_{NPC}}{2}\right)^3 \tag{6}$$

Last, $V_{pn}$ was populated with cellular estimates of NPC species (see Model Addenda and Model Generalization below), which were confined to the perinuclear subcompartment.

Karyopherin-mediated transport through $V_{pn}$ was modeled in the following directional sequence: 1) Cytoplasmic (or nuclear) karyopherin exchanges with the perinuclear subcompartment at the rate used to describe karyopherin-independent nucleocytoplasmic transport in the original model[1]. 2) Perinuclear karyopherin binds to an unoccupied nuclear pore (NUP) according to the kinetic parameters reported in Ref. [2]. 3) The NUP:karyopherin complex unbinds, releasing karyopherin into the nuclear (or cytoplasmic) subcompartment and retaining the dissociated NPC in the perinuclear subcompartment (Fig. SN1). The final model leaves open the possibility of two classes of nuclear pores[10], one that transports all cellular cargo and one that additionally transports a specific cellular cargo. The overall formalism achieves bidirectional transport across the NPC and Identifies when import-export kinetics are limited by the number of available NPCs per cell. In the final model, NPC limitations occur with fewer than ~10 cargo-specific nuclear pores per cell (Fig. SN2). These modeling results are consistent with the enormous cargo-carrying capacity of individual NPCs[11].

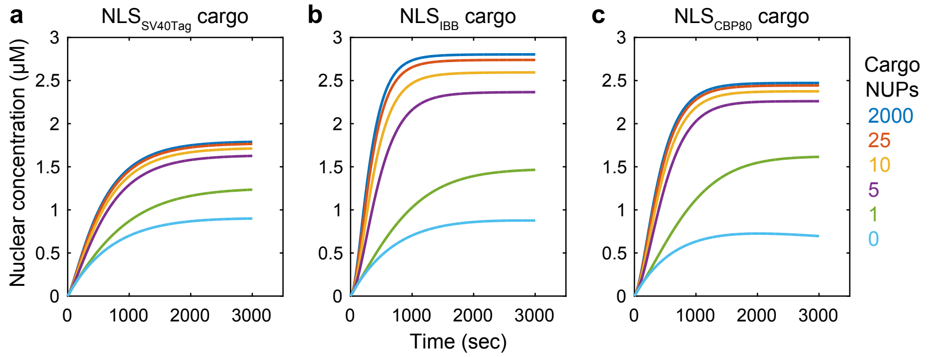


**Fig. SN2.** Evidence for limitations in cargo transport when nuclear pores (NUPs) are extremely limiting. Details on the different NLS cargos are explained later in the text. Simulations were performed by spiking in cargo at a whole-cell concentration of 1 μM.

*Nuclear Export Module*

The nuclear export module of Ref. [2] adds three proteins: a prototypical NES-containing cargo, the exportin CRM1, and the cofactor RanBP3 that enhances CRM1-mediated export[12, 13]. NES cargo, CRM1, and RanBP3 assemble with RanGTP in the nucleus, and the four-protein holocomplex is exported through the NPC according to the transport mechanisms described above. In the cytoplasm, CRM1–RanBP3–NES–RanGTP binds to RanBP1, which enables RanGAP to promote GTP hydrolysis and dissociation of the holocomplex into its individual constituents. RanBP3 contains a conventional NLS, which is reimported into the nucleus via importin-$\alpha/\beta$, and CRM1 passively exchanges between the cytoplasm and nucleus by karyopherin-independent diffusion.

Although Ref. [2] is unclear about the kinetic parameters for recycling RanBP3 and CRM1 to the nucleus and for assembling–disassembling CRM1–RanBP3–NES–RanGTP, we managed to fill voids with existing parameters by making reasonable assumptions. For example, formation of the CRM1 and RanBP3 intermediate complexes with RanGTP were assumed to occur with the same kinetics as the association kinetics listed in Ref. [2] for CRM1–RanBP3 + RanGTP. Likewise, we assumed that the RanBP1-induced disassembly of CRM1–RanBP3–NES–RanGTP was the same as that for RanGTP–CAS–importin-$\alpha$ in Ref. [1] (see Model Corrigenda above). The NLS on RanBP3 was assumed to bind with the same kinetics as $NLS_{SV40Tag}$, and the permeability of CRM1 was assumed to be the same as CAS (another exportin in the model; Supplementary Table 1). The disambiguated nuclear export module thus required no additional rate parameters beyond those specified in or corrected from Refs. [1, 2].

*Model Addenda*

Ref. [2] simulates a bipartite NLS from murine cap-binding protein p80 ($NLS_{CBP80}$), which binds to importin-$\alpha$–importin-$\beta$ with 10-fold higher affinity[14] than the NLS from SV40 T antigen ($NLS_{SV40Tag}$) used in the original model[1]. However, no information is provided about how $NLS_{CBP80}$ was modeled. To retain the two-state binding kinetics of $NLS_{SV40Tag}$ (Eqn. 2), we assumed that $k_{on,1}$ and $k_{on,2}$ of $NLS_{CBP80}$ were identical to $NLS_{SV40Tag}$ (Supplementary Table 1). We then reduced $k_{off,1}$ of $NLS_{CBP80}$ to its lowest order of magnitude (0.01 $s^{-1}$) and fit $k_{off,2}$ to approximate the published affinity ($k_{off,2} = 2.5 \times 10^{-4}$ $s^{-1} \Rightarrow K_{D,eff} = 2.5$ nM ~ 2.4 ± 0.5 nM for $NLS_{CBP80}$[14]). Under these assumptions, import of $NLS_{CBP80}$ cargo is predicted to occur at a faster initial rate and higher steady state compared to $NLS_{SV40Tag}$ cargo (Fig. SN2a,c). The predictions do not fully agree with Ref. [2], likely because of the million-fold correction in cargo affinity described earlier (see Model Corrigenda above). Nevertheless, the transport of $NLS_{CBP80}$ cargo remains slower than the IBB domain of importin-$\alpha$ ($NLS_{IBB}$), a different type of cargo that binds directly to importin-$\beta$. Binding parameters for importin-$\beta$ and $NLS_{IBB}$ were reported in Ref. [1] (with the same $k_{on,2}$ errors as $NLS_{SV40Tag}$; Supplementary Table 1), but they were never used in that work. With the corrected rate parameters, we simulated import of $NLS_{IBB}$ and observed faster initial import and a higher steady state compared to both $NLS_{SV40Tag}$ and $NLS_{CBP80}$ (Fig. SN2). These predictions were qualitatively similar to Ref. [2].

Another omission of Ref. [2] was the reported abundance of "native" NES cargo akin to the native $\alpha$–$\beta$ cargo and native $\beta$ cargo estimated in Ref. [1]. We circumvented this limitation by cross-referencing a database of confirmed NES-containing endogenous proteins[15] with a recent SWATH-MS dataset of absolute protein abundances in HeLa cells[16]. Aggregating the detectable NES-containing proteins yielded an estimate of 1 μM native NES cargo after rounding upwards. We adopted a similar approach to update the prior estimate of native $\beta$ cargo by using two proteomic surveys of $\beta$-binding cargo[17, 18] along with the SWATH-MS resource in HeLa cells[16] to tabulate copy numbers per cell. Aggregating the detectable $\beta$-binding proteins yielded an estimate of 5 μM native $\beta$ cargo after rounding upwards, a fivefold higher estimate than postulated in Ref. [1].

Last, we used the widely cited quantity of 3000 NPCs[19] per HeLa cell as the starting point for defining the initial concentration of nuclear pores in the perinuclear subcompartment. This number is far in excess the NPC counts where transport is predicted to become restricted and thus is not a sensitive initial condition in the model (Fig. SN2). Overall, despite considerable uncertainty in the NPC module of the earlier work[2], our implementation here retains the important characteristics of different NLS cargo types with conservative changes relative to the original modeling framework[1, 2].

*Model Generalization*

The earlier models[1, 2] were developed and tested in HeLa cells; thus, it was important to adapt the new model to B2:B1 cells. We stained trypsinized B2:B1 cells with wheat germ agglutinin and DAPI, segmented, and quantified cells as spheres to arrive at median cytoplasmic and nuclear volumes of $V_{cyt,B2:B1}$ = 1.45 pL and $V_{nuc,B2:B1}$ = 0.52 pL. Our own estimates for HeLa subcellular volumes ($V_{cyt,HeLa}$ = 2.55 pL; $V_{nuc,HeLa}$ = 0.87 pL) were larger than those and used in the Jarnac code of Ref. [1] but agree better with published values[20, 21]. To adapt initial protein concentrations, we performed quantitative immunoblotting[22] on equal numbers of B2:B1 cells and HeLa cells (kindly provided by Dr. Ian Macara, Vanderbilt University) (Fig. SN3). Serial dilutions of lysate established a hyperbolic calibration curve relating arbitrary protein mass (*M*) to integrated band intensity. The differences in arbitrary protein mass were then scaled by the ratio of whole-cell volumes and the reported whole-cell HeLa concentration of the protein ($C_{HeLa}$) from Ref. [1] or Ref. [2]:

$$C_{B2:B1} = C_{HeLa} \frac{M_{B2:B1}/(V_{cyt,B2:B1}+V_{nuc,B2:B1})}{M_{HeLa}/(V_{cyt,HeLa}+V_{nuc,HeLa})} \quad (7)$$

Copy numbers per cell were generally comparable between B2:B1 cells and HeLa cells (Fig. SN3). However, the ~40% smaller volume of B2:B1 cells indicated that whole-cell concentrations of some species were as much as threefold higher (Table SN1). We extended this trend by increasing the estimated concentration of "generic transport receptors" for the HeLa cells[1] (3.6 µM) to 6 µM for B2:B1 cells (Supplementary Table 1).

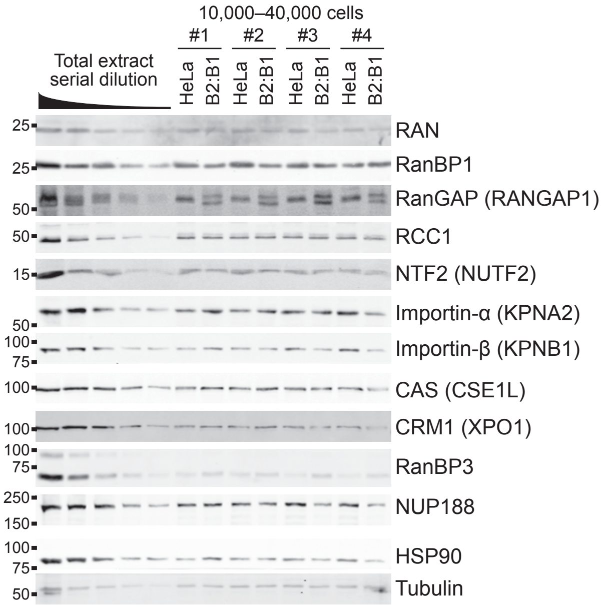


**Fig. SN3.** Quantitative immunoblotting of relative abundances between HeLa cells and B2:B1 cells. Common names from Riddick and Macara[1,2] are listed with the official gene name in parenthesis if different. NUP188 was used as a core nucleoporin to estimate the total number of NPCs, and HSP90 and Tubulin were used as loading controls.

| Model Species | HeLa | B2:B1 | Units |
|---|---|---|---|
| Ran | 5 ± 0.8 | 5.2 ± 0.4 | µM |
| RanBP1 | 2 ± 0.4 | 2.7 ± 0.6 | µM |
| RanGAP | 0.5 ± 0.03 | 0.9 ± 0.2 | µM |
| RCC1 | 0.25 ± 0.01 | 0.3 ± 0.03 | µM |
| NTF2 | 0.6 ± 0.07 | 0.8 ± 0.07 | µM |
| Importin-α | 1 ± 0.2 | 1.5 ± 0.2 | µM |
| Importin-β | 3 ± 0.9 | 2.8 ± 0.3 | µM |
| Cas | 3 ± 0.1 | 5.3 ± 0.9 | µM |
| CRM1 | 0.3 ± 0.02 | 0.5 ± 0.08 | µM |
| RanBP3 | 0.05 ± 0.01 | 0.15 ± 0.01 | µM |
| Nuclear pore complexes | 3000 ± 300 | 2600 ± 500 | copies |

**Table SN1.** Empirically adjusted protein concentrations for B2:B1 cells in the nucleocytoplasmic shuttling model according to per-cell relative copy number estimates (Fig. SN3) and differences in cell volume (Supplementary Table 1).
